# Neurophysiological Correlates of Modifiable Dementia Risk Factors in Cognitively Unimpaired Older Adults

**DOI:** 10.1101/2025.02.20.639395

**Authors:** Jacob M. Levenstein, Ciara Treacy, Sophie C. Andrews

## Abstract

The 2024 Lancet Commission on Dementia estimates that up to 45% of dementia cases could be prevented by addressing modifiable risk factors, emphasising both prevention opportunities and the need to understand the biological mechanisms. This study investigated neurophysiological mechanisms underlying modifiable dementia risk factors in cognitively unimpaired older adults. Seventy-nine cognitively unimpaired older adults underwent MRI brain scans, with spectroscopy measurements taken from the sensorimotor cortex (SMC) and prefrontal cortex (PFC), using a Hadamard Encoding and Reconstruction of MEGA-Edited Spectroscopy (HERMES) sequence, optimised for measuring GABA+. Modifiable dementia risk scores were calculated using the Assessment for Cognitive Health and Dementia Risk (CogDrisk). Hierarchical linear regression analyses revealed a significant negative relationship within the SMC, with lower GABA+ (β = −0.249, p = 0.009) associated with higher risk scores. In the PFC, lower tNAA and tCho concentrations significantly predicted higher risk scores (β = −0.168 and −0.170, respectively). These findings suggest that GABAergic system alterations may underlie the pathophysiology of modifiable dementia risk in healthy ageing, whilst changes in tNAA and tCho may reflect early alterations in neuronal integrity. These region-specific neurochemical findings may help identify potential early biomarkers for dementia risk, and suggest new therapeutic pathways for preventive interventions.

## 1. Introduction

Late-life dementia eventuates as a result of neurodegeneration from chronic disease processes initiated in mid-life (Henstridge et al., 2019) or earlier (Radford et al., 2017), though the driver of these changes remains unclear. The Lancet Commission on Dementia estimates that up to 45% of world-wide dementia cases are attributed to preventable modifiable risk factors (Livingston et al., 2024), and understanding the underlying brain mechanisms driving this modifiable risk is particularly important, given the large potential for prevention. Individualised, validated dementia risk scores are calculated by leveraging recognized protective- and risk-factors of dementia, incorporating into these weighted scores both modifiable and non-modifiable factors (Anstey et al., 2022). Prior investigations of dementia risk scores and structural Magnetic Resonance Imaging (MRI) have highlighted significant relationships with increased accumulation of deep white matter lesions, decreased cortical thickness, decreased white matter, grey matter and hippocampal volumes (Stephen et al., 2017;2021; Heger et al., 2021;Pace et al., 2024). However, the neurophysiology and/or mechanism(s) underlying these observed structural relationships remains unknown.

Proton magnetic resonance spectroscopy (^1^H-MRS) is a non-invasive imaging technique facilitating measures of neurochemical concentrations from human brain tissue. In those living with mild cognitive impairment (MCI) and dementia, N-acetylaspartate (NAA), a marker of neuronal integrity/loss and/or demyelination, Myo-Inositol (Myo-Ins), a marker of glial activation and inflammation, and Choline (Cho), a marker of neurodegeneration and/or inflammation, have been most commonly studied with some studies demonstrating reduced levels in comparison to healthy older adults (Rae 2014; McKiernan et al., 2023). In contrast, gamma aminobutyric acid (GABA), the primary inhibitory neurotransmitter and key regulator in neural oscillatory dynamics, is less commonly studied in the context of dementia (McKiernan et al., 2023). This is in part due to technical requirement of metabolite-tailored edited-MRS sequences to resolve overlapping spectral profiles (Saleh et al., 2016), and partially due the notion of GABAergic resilience in AD (Rossor et al., 1982), such that GABAergic neurons are more resistant to beta amyloid plaques than glutamatergic or cholinergic neurons (Pike et al., 1993; Xu et al., 2020), supporting a narrative that GABAergic neurons were not involved in the disease progression. However, recent studies have begun suggesting a larger role for GABA in dementia. For example, GABAergic hypoactivation may drive excitation/inhibition (E:I) imbalance, which in turn potentiates pathology (Govindpani et al., 2017, Carello-Collar et al., 2023), particularly early in the disease process, such as in pre-clinical or higher risk individuals (Bi et al., 2020). Furthermore, findings of GABAergic inhibitory interneuron deficits in AD identify decreased activity of GABA interneurons and synaptic release increasing aberrant neural network activity and contributing to cognitive deficits (Xu et al., 2020). However, to date, it remains untested if GABA concentrations are associated with modifiable dementia risk severity.

To address this question, we investigated the underlying neurophysiology of modifiable risk factors of dementia in a cohort of healthy older adults. We employed a multi-edited ^1^H-MRS sequence to measure GABA concentrations from two cortical regions, the sensorimotor cortex (SMC) and prefrontal cortex (PFC). Our primary aim was to investigate if GABA from SMC and/or PFC are significantly associated with modifiable dementia risk scores in an ageing population, absent of a dementia diagnosis or cognitive impairment. We hypothesise that healthy ageing individuals with lower GABA concentrations will have greater modifiable dementia risk scores. The second aim was to investigate whether alternative neurochemicals implicated in prior case-control dementia studies were significant predictors of modifiable dementia risk scores. These secondary neurochemicals include: total NAA (tNAA), Myo-Ins, total Cho (tCho), combined glutamate and glutamine (Glx), E:I and Glutathione (GSH). We hypothesise that healthy ageing individuals with greater modifiable dementia risk scores will have lower tNAA, higher Myo-Ins, lower tCho, lower Glx, greater E:I imbalance and lower GSH. Finally, a post-hoc analysis examining neurochemical regional differences between the SMC and PFC are reported to aid in the interpretation of the findings.

## 2. Methods

### 2.1 Participants

This study was approved and conducted in adherence with the Human Research Ethics Committee of the University of the Sunshine Coast (S211620), which adheres to the Declaration of Helsinki and the Australian National Statement on Ethical Conduct in Human Research (2023). All participants provided informed written consent prior to commencement of study participation. Right hand dominant healthy individuals aged 50 – 85 were screened for study eligibility based on the following criteria: no history or diagnosis of a neurological, cerebrovascular, respiratory, or psychological condition, no prior head injury, uncontrolled diabetes, medications impacting the central nervous system, current substance abuse or misuse, and free from MRI contraindications. A total of 82 participants enrolled in the study. Two participants were excluded due to unreported MRI contraindications during prescreening. One participant was removed due to poor comprehension of the study instructions resulting in unusable data. The final cohort included 79 participants with an age range of 50.6 to 84.8 years old. The mean age was 68.1 years (SD = 9.65), and the gender distribution was fairly balanced, with 44 female and 35 male participants included. The education level ranged from 9 to 26 years, with a mean of 16.12 years (SD = 3.76).

### 2.2 Modifiable Dementia Risk Scores

Dementia risk scores were generated from the Assessment for Cognitive Health and Dementia Risk (CogDrisk; Anstey et al., 2022), providing validated individualised dementia risk scores (Kootar et al., 2023). The CogDrisk appraises 17 modifiable and non-modifiable risk factors of dementia, weighting each factor based on current evidence (Figure 1.A). A recent comparison of the CogDrisk and other validated dementia risk tools (Australian National University–Alzheimer Disease Risk Index (ANU-ADRI; Anstey et al., 2014), Cardiovascular Risk Factors, Aging and Dementia (CAIDE; Kivipelto et al., 2006, and Lifestyle for Brain Health (LIBRA; Schiepers et al., 2018) concluded that the CogDrisk performed similarly, whilst also including a greater number of risk factors and more up-to-date factor weightings (Kootar et al., 2023). By design, the exclusion criteria of this study screened out potential participants with known non-modifiable risk factors, including a history of traumatic brain injury, stroke, or uncontrolled diabetes. After adjusting dementia risk scores for the remaining non-modifiable risk factors (age, education, and gender) the resultant dementia risk scores reflect only the modifiable risk factors, such as high cholesterol, hypertension, obesity, arterial fibrillation, insomnia, depression, physical activity, cognitive engagement, social engagement, diet and smoking. The Dementia Risk Score was available for 78 participants, ranging from −1.25 to 24 (Figure 1.B), with a mean of 10.54 (SD = 6.11). The wide range and relatively large standard deviation indicate considerable variation in dementia risk among participants.

**Figure 1:**
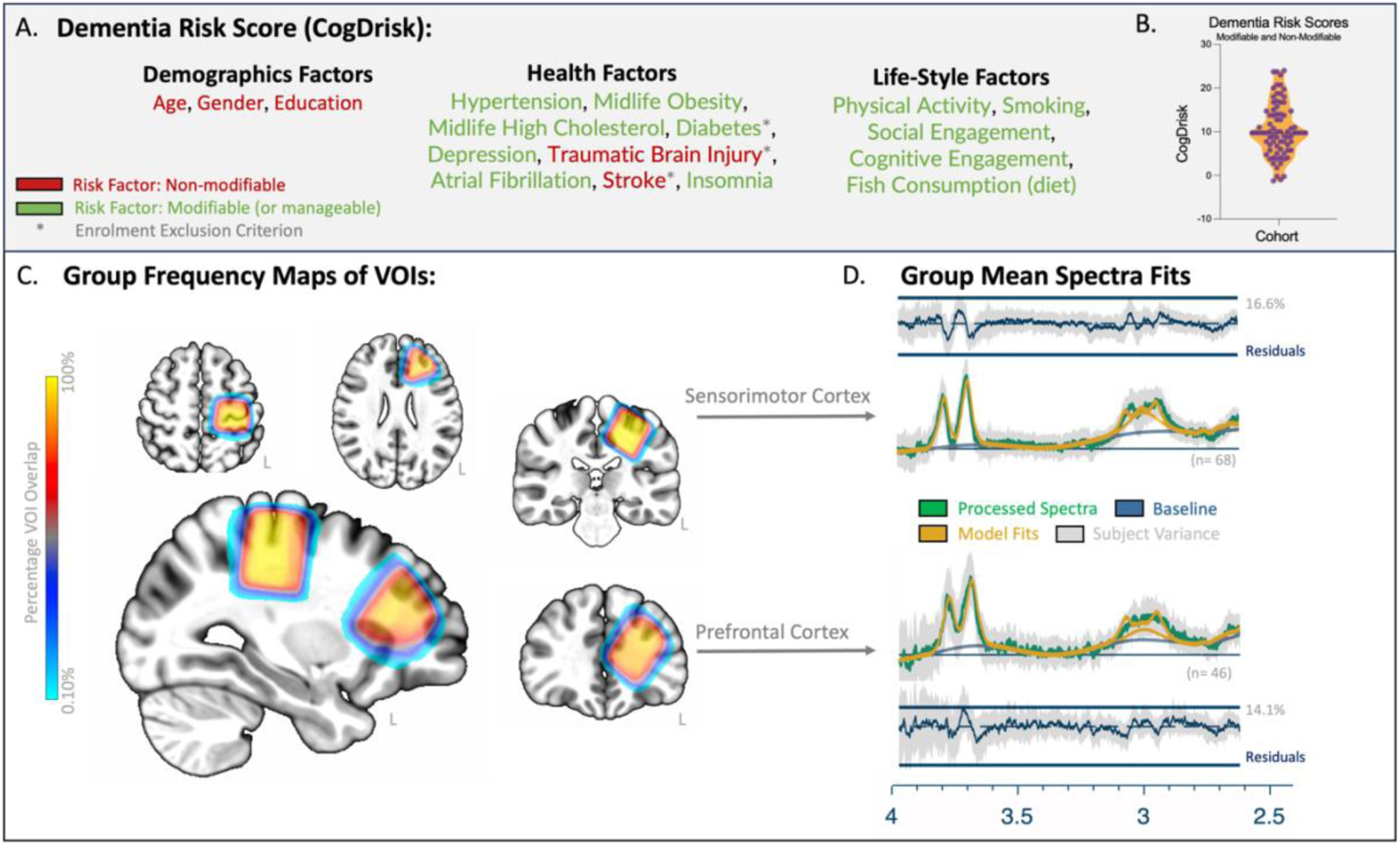
A. Visualization of dementia risk score factors denoted as non-modifiable (red) and modifiable (green). B. Distribution plot of dementia risk scores across the cohort. C. Frequency plot depicting the percentage of spatial overlap of participants’ sensorimotor cortex and prefrontal cortex spectroscopy voxels in standard space. D. Group mean spectroscopy data (green) with overlaid model fits (yellow) for GABA and Glx, residuals of model fits (dark blue) and SEM across participants (light gray).

As a screen for cognitive impairments, participants completed the the Montreal Cognitive Assessment (MoCA; Nasreddine et al., 2005). 71 participants completed the MoCA, with scores ranging from 23 to 30, with a mean of 27.31 (SD = 1.86). As the MOCA’s maximum score is 30, and scores above 26 are typically considered normal, these results suggest that most participants had relatively intact cognitive function, an no participants’ MoCA scores were within the range of moderate or severe cognitive impairments (i.e., scores below 17).

### 2.3 Magnetic Resonance Imaging Acquisition and Processing

All participants completed an MRI brain scan at the Nola Thompson Centre for Advanced Imaging (Thompson Institute, University of the Sunshine Coast), via a 3T Skyra (Siemens, Erlangen Germany) and 64 channel head-coil. The MRI protocol consisted of a T1-weighthed magnetization prepared rapid acquisition gradient echo sequence (MPRAGE: TR = 2200ms, TE = 1.71ms, TI = 850ms, flip-angle = 7°, spatial resolution = 1mm^3^, FOV = 208×256x256,TA = 3:57). This scan was used to plan the placement of the two MRS volume of interest (VOI) voxels. Two single-voxel MR spectroscopy scans were acquired from the left SMC and left PFC, each using a Hadamard Encoding and Reconstruction of MEGA-Edited Spectroscopy (HERMES) sequence (Chan et al., 2016), optimised for measuring GABA and GSH (Saleh et al., 2016); VOI=3cm^3^, TR=2000ms, TE=80ms, flip-angle=90◦, averages=320, edited pulse 1 = 1.9pmm, edited pulse 2 = 4.56ppm, edited off pulse = 7.5ppm, TA=10:48). With respect to the sequence required VOI size and the age of the study population, the left SMC and left PFC were both selected as a cortical sites, providing enough cortical tissue for VOI placement, avoiding subcortical regions, lateral ventricles and dura.

All participants’ T1-weighted scans were visually inspected for image quality, and (re)confirming accuracy of VOI placement (figure 1.C). Zero T1-weighted scans were removed due to poor data quality or poor VOI placement. MRS data was analysed using OSPREY’s processing and fitting pipeline (v.2.4.0; Oeltzschner et al., 2020). All MRS data and experiment conditions were visually inspected to verify data quality and accuracy of model fits (see figure 1.D.) As all neurochemicals are presented as a ratio over creatine (Cr) and phosphocreatine (PCr; combined as tCr), excluded sum experiments based on poor tCr fits resulted in a listwise exclusion (i.e., excluded sum, diff1 and diff2).

From the sum experiments, poor data quality impacting model fits for tCr, NAA, NAAG (combined as tNAA), Myo-Ins, glycerophosphocholine and phosphocholine (combined at tCho) resulted in listwise exclusion of seven datasets from SMA and twelve datasets from PFC. From the diff1 experiments, poor data quality impacting the model fits for GABA and MM3co (combined as GABA+) and/or Glx resulting in case-wise exclusion of three GABA+ and Glx datasets from SMA and twenty-one GABA+ and twelve Glx from PFC. From Diff2 experiments, poor data quality impacting the model fits for GSH resulting in case-wise exclusion of five dataset for SMA. Finally, one participant had not completed the SMC scan, resulting in a max SMC cohort size of 71 and a max PFC cohort size of 67.

### 2.4 Statistical Analyses and Assumption testing

All statistical analyses were performed using SPSS (v.29.0, IBM Corp, 2023). Variables were reviewed for normality, with outlier correction based on a z-score standard deviation transformation method (i.e., z-scores data values >3.29 or <3.29 were adjusted to one unit above or below the nearest value existing within the acceptable ranges; Tabachnick et al., 2013). Three measures contained a single outlier value: SMC-GABA+, SMC-E:I and PFC-GSH. Shapiro-Wilk’s tests of normality was performed for all measures of interest (i.e., age, education, dementia risk scores and seven neurochemicals per VOI). Only Age (W = 954, *p* = 0.006), education (W=0.969, *p* = 0.047), and SMC-E:I (W=0.768, p < 0.001) were not normally distributed. Prior to testing the primary and secondary aims, bivariate correlations were undertaken with age, and the seven neurochemicals of interest across the two MRS VOIs. These bivariate correlations were repeated for Education and dementia risk scores with the seven neurochemicals of interest. Spearman’s rank (Rho) tests were used for any relationship tested containing SMC-GABA+, SMC-E:I or PFC-GSH. Pearson’s r was performed for all remaining bivariate relationships containing normally distributed variables.

For the primary and secondary aims, hierarchical linear regression was performed independently for each of the seven neurochemicals of interest across the two regions of interest. All independent regression models contained age, gender and education entered in step 1, with step 2 containing single neurochemical regressor of interest. Post-hoc analyses of regional differences in neurochemical concentrations from SMC and PFC were conducted to determine amplitude differences by employing paired t-tests or Wilcoxon Signed Rank Tests (with respect to the normality of the dataset) and across cohort rank differences employing Spearman’s rank correlations.

## 3. Results

### 3.1 Bivariate Relationships of Demographics, Neurochemicals and Dementia Risk Scores

For Age, bivariate correlations revealed no significant relationships with neurochemicals from SMC (*p.s.* => 0.263) or for PFC (*p.s.* => 0.061). For education, no significant relationships with neurochemicals were identified for SMC (*p.s.* => 0.286) or for PFC (*p.s.* => 0.109). For dementia risk scores (containing age, gender and education), SMC-GABA+ was significantly correlated (*r* = −0.725, *p* = 0.008), such that individuals with lower GABA+ concentrations had greater dementia risk. The remaining six SMC relationships were not significantly correlated with dementia risk scores (*p.s.* => 0.053). For PFC, tNAA (*r* = −0.243, *p* = 0.049) and PFC-tCho (r = - 0.246, *p* = 0.046) were each significantly correlated with dementia risk scores, such that individuals with lower tNAA and lower tCho had greater dementia risk scores. The remaining five PFC relationships were not significantly correlated with dementia risk scores (*p.s.* => 0.251).

### 3.2 Sensorimotor Cortex Neurochemical Predictors of Modifiable Dementia Risk

Results from hierarchical linear regression revealed that SMC-GABA+ was significantly associated with modifiable dementia risk scores, within a model containing age, gender, education and SMC-GABA+ (F(4,63) = 16.73, p < 0.001, R^2^ = 0.515). The addition of SMC-GABA+ to the step one model (F(3,64) = 18.16, *p* < 0.001, R^2^ = 0.460) resulted in a significant improvement (FΔ = 7.170, Sig. FΔ = 0.009). After controlling for non-modifiable risk-factors, GABA+ significantly modelled modifiable dementia risk (β = −0.249, *t* = −2.68, *p* = 0.009, ΔR^2^ = 0.055), such that individuals with lower SMC-GABA+ had greater dementia risk scores (figure 2.A.1).

**Figure 2:**
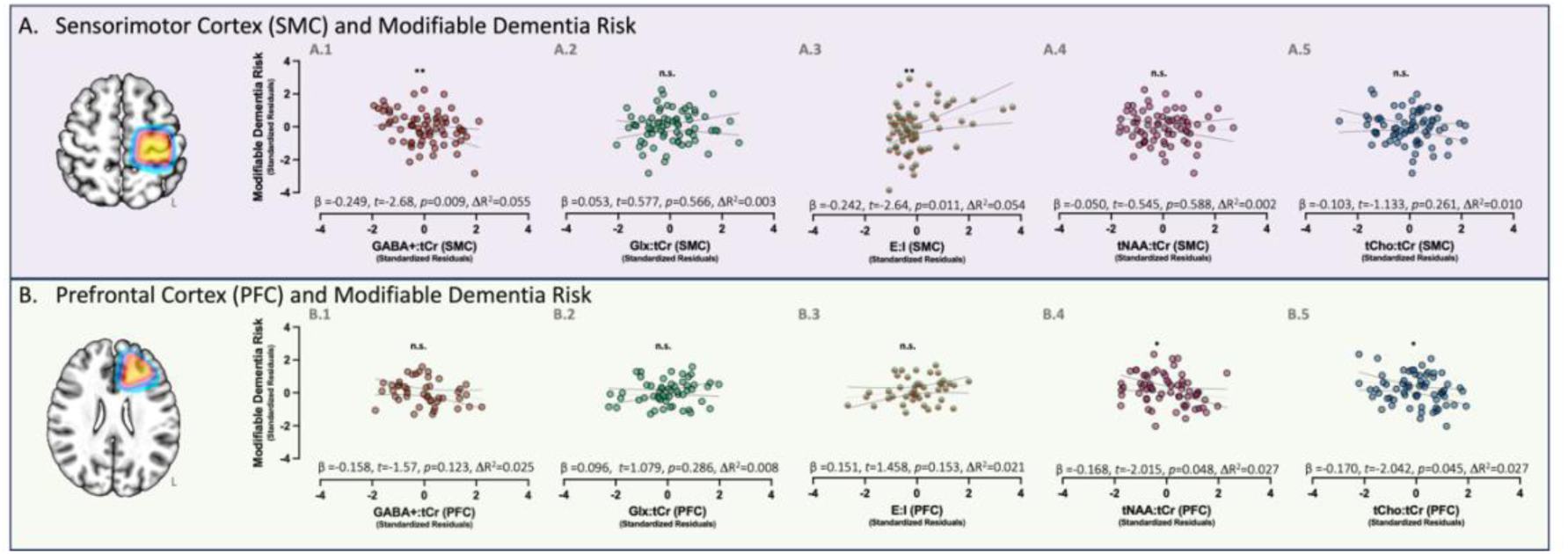
Hierarchical linear regression partial correlation plots of neurochemical concentrations (x-axis) and Modifiable Dementia Risk Scores (y-axis), after controlling for age and education. Top row (A) depicts relationships using neurochemical concentrations from left sensorimotor cortex. Bottom row (B) depicts relationships using neurochemical concentrations from left prefrontal cortex. \**p* < 0.05, \*\**p* < 0.01. Light-grey dotted lines depict linear regression fit, dark-grey dotted lines depict upper and lower 95% confidence intervals.

Next, secondary neurochemicals from SMC were tested via six independently conducted hierarchical linear regression models, one per neurochemical of interest (i.e., tNAA, Myo-Ins, tCho, Glx, E:I and GSH). Of the six models tested, only SMC-E:I was identified as a significant neurochemical predictors of modifiable dementia risk, via a model containing age, gender, education and E:I (F(4,63) = 16.62, *p* < 0.001, R^2^=0.513). The addition of SMC-E:I to this model resulted in a significant improvement (FΔ = 6.948, Sig. FΔ = 0.011). After controlling for non-modifiable risk-factors, SMC-E:I was significantly associated with modifiable dementia risk (β = 0.242, t = 2.64, *p* = 0.011, ΔR^2^=0.054), such that individuals with greater SMC-E:I imbalance had greater dementia risk scores (figure 2.A.3). For the remaining five models, none significantly improve the model of modifiable dementia risk scores above what was explained by age, gender and education (*p.s*. => 0.261).

### 3.3 Prefrontal Cortex Neurochemical Predictors of Modifiable Dementia Risk

Results from hierarchical linear regression revealed that the addition of PFC-GABA+ to the model with age, gender and education did not result in a significant improvement (FΔ = 2.476, Sig. FΔ = 0.123) from step 1. Next, secondary PFC neurochemicals were investigated. Two of the six independent models were identified as significant neurochemical predictors of modifiable dementia risk. First, PFC-tNAA was significantly associated with modifiable dementia risk scores from a model containing age, gender, education and PFC-tNAA (F(4,61) = 22.60, *p* < 0.001, R^2^ = 0.597). The addition of PFC-tNAA to this model resulted in a significant improvement (FΔ = 4.061, Sig. FΔ = 0.048). After controlling for non-modifiable risk-factors, PFC-tNAA significantly modelled modifiable dementia risk (β = −0.168, *t* = −2.02, *p* = 0.048, ΔR^2^ = 0.027), such that individuals with lower PFC-tNAA had greater dementia risk scores (figure 2.B.4). Second, PFC-tCho was significantly associated with modifiable dementia risk scores via a model containing age, gender, education and PFC-tCho (F(4,61) = 22.66, *p* < 0.001, R^2^ = 0.598). The addition of PFC-tCho to this model resulted in a significant improvement (FΔ = 4.170, Sig. FΔ = 0.045). After controlling for non-modifiable risk-factors, PFC-tCho significantly modelled modifiable dementia risk (β = −0.170, *t* = −2.04, *p* = 0.045, ΔR^2^ = 0.027), such that individuals with lower PFC-tCho had greater dementia risk scores (figure 2.B.5). For the remaining four models (analysed independently), none significantly improve the model of modifiable dementia risk scores above what was explained by age, gender and education (*p.s*. => 0.123).

### 3.4 Regional Differences in Neurochemicals

Of the seven neurochemicals tested, all but GABA+ and GSH had significant amplitude differences between SMC and PFC concentrations (*p.s.* < 0.001, see table 1). For tNAA and E:I, values from SMC were significantly greater than PFC. For Myo-Ins, tCho and Glx, values from SMC were significantly lower than PFC. With respect to rank relationships of neurochemical concentrations from SMC and PFC, tNAA and tCho each significantly rank-correlated in their measures from SMC and PFC measures (*p.s.* < 0.001), with the remaining five neurochemicals not having significant rank differences (*p.s.* >= 0.153).

**Table 1:** Paired Samples T-Test.

| Measure 1<br>SMC | Measure 2<br>PFC | t | df | p | Mean Difference | SE<br>Difference | Cohen's d | SE Cohen's d |
| --- | --- | --- | --- | --- | --- | --- | --- | --- |
| tNAA | - tNAA | 9.817 | 60 | 4.276×10<sup>-14</sup> | 0.272 | 0.028 | 1.257 | 0.185 |
| Myo-Ins | - Myo-Ins | -6.351 | 60 | 3.165×10<sup>-8</sup> | -0.150 | 0.024 | -0.813 | 0.189 |
| tCho | - tCho | -18.269 | 60 | 3.418×10<sup>-26</sup> | -0.052 | 0.003 | -2.339 | 0.160 |
| Glx | - Glx | -4.677 | 50 | 2.246×10<sup>-5</sup> | -0.267 | 0.057 | -0.655 | 0.203 |
| GABA+ | - GABA+ | 0.676 | 37 | 0.503 | 0.023 | 0.034 | 0.110 | 0.213 |

|  |  | W | z | p | Hodges-Lehmann<br>Estimate | Rank-Biserial<br>Correlation | SE Rank-Biserial<br>Correlation |
| --- | --- | --- | --- | --- | --- | --- | --- |
| GSH | - GSH* | 879.0 | 0.417 | 0.679 | 0.007 | 0.064 | 0.151 |

\*Wilcoxon signed-rank test. Unless otherwise denoted, all tests were Student's t-tests

## 4. Discussion

This study provides novel insights into the neurophysiological markers of modifiable dementia risk in healthy ageing adults. The primary aim of the study was to investigate whether MRS-derived GABA+ concentrations related to individuals’ modifiable dementia risk scores. As hypothesised, GABA+ was identified as a significant predictor of modifiable dementia risk scores, such that individuals with greater dementia risk had lower GABA+. This finding was region specific, only identified from concentrations within SMC, and not within PFC. The secondary aim - investigating six alternative neurochemicals - identified SMC-E:I, PFC-tNAA and PFC-tCho were each significant predictors of modifiable dementia risk scores, such that individuals with greater modifiable dementia risk had greater SMC-E:I imbalance, lower PFC-tNAA and lower PFC-tCho. The remaining neurochemical null findings (i.e., Myo-Ins, Glx, GSH) suggest that previously identified neurochemical differences observed in mild cognitive impairment and dementia, may not be present in-relation to modifiable dementia risk scores collected from a cognitively unimpaired ageing cohort. These novel results identify potential early indicators of spatial and neurochemically distinct changes in relation to modifiable dementia risk.

### 4.1 GABA+ and E:I Imbalance Significantly Associated with Modifiable Dementia Risk

Our finding of lower GABA+ concentrations with increased dementia risk provides the first evidence linking GABAergic system function to modifiable risk factors in healthy ageing. As such, the significant findings reported here within a healthy ageing population provides unique evidence to the growing role of GABA in dementia pathology (McKiernan et al., 2023; Xu et al., 2020;Pike et al., 1993; Govindpani et al., 2017; Carello-Collar et al., 2023; Bi et al., 2020; Li et al., 2016). The significant SMC-GABA+ relationship with modifiable dementia risk may represent an early biomarker, demonstrating how life-style factors impact neurophysiology, which could intern increase or decrease future disease onset. Further, E:I balance disruptions have also been identified as having a detrimental role in AD pathogenesis (Bi et al., 2020), preceding the onset of clinical symptoms by some decades (Palop et al., 2016;Long et al, 2019). The significant SMC-E:I imbalance relationship identified here, in combination with the absence of SMC-Glx as a significant predictor, suggests further the interaction of GABA+ and modifiable dementia risk. As suggested elsewhere, mediating GABA could represent a viable therapeutic target (Calvo-Flores Guzmán et al., 2018) for restoring the E:I balance.

### 4.2 tNAA and tCho Significantly Associated with Modifiable Dementia Risk

The significant associations between prefrontal cortex tNAA and tCho concentrations and modifiable dementia risk extend previous findings from clinical populations to healthy ageing adults (see review: McKiernan et al., 2023). While these relationships explained a smaller portion of variance compared to the SMA GABA+ findings, their presence in cognitively unimpaired individuals suggests they may be sensitive early markers of neural health. The prefrontal-specific nature of these findings aligns with known patterns of cortical vulnerability in early neurodegeneration and may reflect region-specific sensitivity to modifiable risk factors.

### 4.3 Neurochemicals not significantly associated with Modifiable Dementia Risk

The absence of significant relationships between modifiable dementia risk and Myo-Ins, Glx, and GSH provides important context for understanding the progression of neurochemical changes in relation to modifiable dementia risk. These null findings in our healthy ageing cohort, contrasting with previous findings in clinical populations, suggest that alterations in these markers may emerge later in the pathological process. This temporal specificity of neurochemical changes could help establish a biochemical timeline of modifiable dementia risk progression. Alternatively, our study consisted of a single healthy ageing cohort, and tested metrics of interest as a continuum. As such, these null findings may not be comparable with prior clinical studies, which performed group differences between individuals with a diagnosis and matched controls.

### 4.4 Cortical Region Differences in Neurochemicals

GABA+ and GSH were not significantly different in amplitude between SMC and PFC, as both neurochemical concentrations are much lower in concentration than the other neurochemicals, reducing the numerical range and thus sensitivity to detect difference. Further, the non-significant relative rank correlation between GABA+ from SMC and PFC suggest distinct regional information. tNAA and tCho each significantly correlating in rank between the two cortical regions, yet only MRS measures from PFC significantly relating to modifiable dementia risk. As the sample-size differences between regions was unlikely to influence the sensitivity of the tests (SMC-n = 71, and PFC-n = 67), these region-specific relationships suggests that the relative amplitude differences between participants, and not pure rank, is what drove these identified PFC relationship and a factor to consider in the null findings for SMC. Finally, with respect to the regional specificity of GABA+ and E:I, these results may reflect differences in data quality. PFC containing 46 QC passed GABA+ datasets compared to 68 QC passed GABA+ datasets in SMC. Thus, SMC had greater statistical power than PFC and this may have contributed to the PFC-GABA+ null result.

### 4.5 Limitations

While there is clear and abundant need for multidomain interventions targeting modifiable dementia risk reduction as a prevention approach (Treacy et al., 2023, Livingston et al., 2024), this complementary investigation of modifiable dementia risk is cross-sectional, and unable to determine how the identified relationships with neurophysiology may relate to future cognitive decline or reserve. Future studies should measure if interventions targeting modifiable dementia risk reduction may correspond to neurochemical changes, and/or if baseline neurochemical concentrations are predictive of intervention outcomes and/or future diagnosis.

Furthermore, we investigate only the modifiable components of dementia risk and by design, exclude or control for non-modifiable risk factors of dementia. As such, it may be that certain neurochemicals are more related to the non-modifiable risk factors than those that are modifiable. Finally, within the cohort tested, the modifiable dementia risk scores were generally low overall, so it remains unknown if higher modifiable dementia risk scores may change the strength of these relationships.

Finally, with respect to diversity, equity and inclusion, the study recruitment was conducted as a rolling admission of community dwelling individuals. As such, the study population was representative of the local area, which is primarily white, of European ancestry, and of middle to high socioeconomic status. The results should be viewed in the context of the population sampled.

## 5. Conclusion

Our results demonstrate that within the sensorimotor cortex, lower GABA+ and greater E:I imbalance were significantly associated with greater modifiable dementia risk scores, suggesting that the GABAergic system may underlie the pathophysiology of modifiable dementia risk in healthy ageing. Within the prefrontal cortex, lower tNAA and lower tCho were also significantly associated with greater modifiable dementia risk scores, which may reflect early neuronal integrity and/or neuronal density changes relating to modifiable dementia risk. The distinct regional patterns of neurochemical associations with dementia risk highlight the importance of multi-region investigations in understanding brain health, and might reflect distinct mechanisms through which lifestyle factors may influence brain health. These findings establish potential biomarkers for early detection of dementia risk and suggest new therapeutic pathways for preventive interventions.

## ACKNOWLEDGMENTS

We would like to extend our gratitude to the participants for their time and contribution to science.

## CONFLICT OF INTEREST STATEMENT

All authors declare no competing interests, or conflicts of interests.

## FUNDING SOURCES

This work was supported by a SPARK Grant (0980028321), University of the Sunshine Coast

## DATA AVAILABILITY STATEMENT

The datasets presented within this study are available from the corresponding author on reasonable request.

## AUTHOR CONTRIBUTIONS

Dr Jacob M. Levenstein: Conceptualization, Methodology, Software, Formal Analysis, Investigation, Data Curation, Writing – Original Draft, Review and Editing, Visualizations, Supervision, Project Administration, Funding Acquisition. Ms. Ciara Treacy: Conceptualization, Methodology, Investigation, Data Curation, Project Administration, Writing – Review and Editing. Dr Sophie C. Andrews: Conceptualization, Methodology, Project Administration, Writing – Review and Editing

## Notes

### Competing Interest Statement

The authors have declared no competing interest.

